# ACNN-6mA Prediction of N6-Methyladenine Loci in Multiple Species Based on Rice Dataset Pre-training Model^†^

**DOI:** 10.1101/2022.11.14.516303

**Authors:** JianGuo Bai, Hai Yang

**Author notes:** Electronic Supplementary Information (ESI) available: [details of any supplementary information available should be included here]. See DOI: 10.21203/rs.3.rs-1997163/v1.

## Abstract

N6-methyladenine is an epigenetic modification that plays a significant role in various cellular processes. Genome-wide monitoring of methylation sites is conducive to understanding the biological function of methylation. Due to the limitations of traditional dry and wet experiments, a series of machine learning and deep learning methods have been developed to detect methylation sites, but their detection species are single or performance is poor. First of all, we conducted sufficient experiments on the widely studied rice datasets, and compared with the previous research, we have greatly improved in various indicators on the two rice datasets. Then we used the models trained on the rice dataset to fine-tune training in half of the other 11 datasets and predict the other half of the independent datasets. Then we used 11 trained models to test 11 species respectively. It was found that ACNN-6mA could obtain higher AUC, ACC and MCC whether cross-species prediction or independent verification set prediction. ACNN-6mA model and code for follow-up researchers is provided as an open-source tool available at https://github.com/jrebai/ACNN-6mA.

## 1 Introduction

Epigenetic modification is a key modification that regulates gene expression without changing the DNA sequence, and methylation is an extensive epigenetic modification studied in different species’ genomes. N6-methyladenine(6mA), 5-hydroxymethylcytosine (5hmC), and N4 methylcytidine (4mC) are several methylation types of concern. 4mC is catalyzed by N-4 cytosine-specific DNA methyltransferase (DNMT), which can methylate the fourth amino group of cytosine in DNA (Timinskas et al., 1995). 4mC modifications play a considerable role in DNA replication and error correction, cell-cycle function, regulation of gene expression levels, and self-and non-self DNA differentiation, and are involved in genome stabilization, recombination, and evolution^1–4^. 5 hmC is produced from 5mC by the Ten eleven translation (Tet) family protein. 5hmC can combine different regulatory elements to regulate gene expression and maintain pluripotency, nervous system development, and tumorigenesis of embryonic stem cells (ESCs)^5,6^. 6mA is involved in a wide range of biological processes in prokaryotes, playing a prominent role in DNA replication, DNA repair, transcription, gene expression regulation, chromatin conformational remodeling, cell defense, etc^7–13^. Although the specific function of m6A in eukaryotes is not completely determined, some researches have found that it has similar characteristics with prokaryotes^14–16^. In the mid-1950s, 6mA was found in Escherichia coli through traditional technology^17,18^. However, due to inconsistent detection results, mature technology for detecting 6mA was not available until 2015^19,20^. Due to the technical limitations of traditional dry and wet experiments in detecting methylation sites, it is a long-term feasible and promising direction to predict methylation sites using computational methods.

Up to now, some models using machine learning and deep learning to detect 4mC, 5hmC and 6mA sites have been proposed based on different coding methods. MM-6mAPred^21^ regards nucleotide sequence as a Markov chain and uses the Markovmodel-based method on the 6mA-rice-Chen^22^ dataset to identify the 6mA site by using transfer probability between adjacent nucleotides. IDNA-6mA Rice^23^ proposed a rice 6mA dataset with 154000 positive samples and 154000 negative samples respectively. As a representative 6mA site recognition model, it uses the one-hot preprocessing method and Random Forest machine learning algorithm for classification. I6mA-Fuse^24^ constructs five Random Forest models using five separate feature encoding methods: one-hot, di-nucleotide binary, k-space spectral nucleus, k-mer, and EIIPs, and uses linear regression models to combine the prediction probability scores of five RF models based on a single encoding to predict the m6A sites of F. vesca and R. chinensis. I6mA-CNN^25^ uses one-hot, dinucleotide binary, dinucleotide property and a hybrid encoding method to encode the sequence, and then conducts convolution and full join operations to obtain four models and perform fusion to predict methylation sites. Deep6mA ^26^ encodes the sequence as one pot, extracts, and abstracts the features of the sequence using a 5-layer convolutional neural network, and then uses LSTM to model the extracted abstract features in terms of time series, extracts the timing effect between high-level features, and then makes a full connection prediction.

Compared with 4mC^27–37^ and 6mA^22,38–46^ methylation site prediction, there are few studies related to 5hmC site prediction^47,48^, but the basic research method is to train the model based on feature selection. According to the above research, we found that most models based on single methylation site prediction have room for improvement in performance, and most studies are relative to one or two species for site identification. Moreover, it has been found^49,50^ that the prediction of methylation sites by most models is limited to a few species, resulting in poor prediction performance for other species. Therefore, there is an urgent need for models capable of cross-species prediction or prediction on multiple species. We propose a pretraining architecture on the rice dataset, which can be trained on more than 6mA datasets of different species, and has a great improvement in various indicators compared with the latest model on the independent test set.

## 2 Methods

### 2.1 Data and preprocessing

As shown in Table 1, in this study, our datasets are divided into two parts: one is rice datasets based on two datasets^22,23^, which are respectively used as a 10-fold cross-validation method and independent validation set; The other part is a 6mA multi-species dataset^48^. The data of these two datasets are more reliable and evenly distributed. It is convenient to compare our method with previous methods through these public datasets. Among them, lv’s ^23^ rice dataset was used as 10-fold cross-validation data, and the 6mA-rice-Chen^22^ dataset was directly tested on the trained dataset without any training. For the rice dataset and the multispecies dataset, we divided the samples of each species into two parts with the same positive and negative ratio, one as the finetuning training set based on the rice model, and the other as the independent validation set.

**Table 1.** Details of datasets used in this experiment.

| species | train | test |
| --- | --- | --- |
| A. thaliana | 15937 | 15936 |
| C. elegans | 3981 | 3980 |
| C. equisetifolia | 3033 | 3033 |
| D. melanogaster | 5596 | 5595 |
| F. vesca | 1551 | 1551 |
| H. sapiens | 9168 | 9167 |
| R. chinensis | 300 | 300 |
| S. cerevisiae | 1893 | 1893 |
| Tolypocladium | 1690 | 1689 |
| T. thermophile | 53800 | 53800 |
| Xoc. BLS256 | 8608 | 8607 |
| rice-lv | 154000 | 154000 |
| rice-chen | 880 | 880 |

Some basic biological and chemical features, Research evidence, of nucleic acids stand behind the frequencies of dinucleotides^51^. Therefore, compared with using one-hot coding directly, di-nucleotide binary coding is able to obtain more information related to DNA structure, which makes it easier to extract sequence information. For ATCG, 16 combinations can be formed (AA AT AG AC TA TT TG TC GA GT GG GC CA CT CG CC). For DNA sequence data, we use the dinucleotide binary encoding method to process sequence data. For example, for “ATCGATGTAC”, we can get combinations (AT TC CG GA AT TG GT TA AC). For each combination, one-hot preprocessing is used. Since the data we obtained are all 41bp, one-hot preprocessing is used to code this combination, and each sequence is coded as a matrix of 40×16.

### 2.2 ACNN-6mA framework

Figure 1 describes the network structure and flow of ACNN-6mA. ACNN-6mA first takes the dinucleotide binary code of the sequence as the input and then carries out feature extraction and automatic combination between advanced features through five attraction-CNN modules to predict methylation sites. The submodules of each attention-CNN module can be added and disassembled freely. They are mainly the following eight submodules: one-dimensional convolution layer, one-dimensional Pooling layer, BatchNormalization layer, Dropout layer, Channel shuffle^52^ layer, one-dimensional ECA^53^ (efficient channel attachment) layer, one-dimensional SA^54^ (spatial attachment) layer, and full connection layer. The Pooling layer and Dropout layer are regularization measures, which can effectively prevent overfitting the network too quickly. And because they can greatly reduce the network parameters, also accelerate the fitting speed. We use the channel shuffle layer to disrupt the channels of features extracted from each layer, which can prevent overfitting and learning more features. We creatively transformed the ECA module and SA module in image processing into a one-dimensional ECA layer and SA layer and made them available as separate modules. Using the ECA layer and SA layer can make the prediction of methylation sites more accurate by applying different attention to different channels and specific sites in different sequences.

**Fig. 1.**
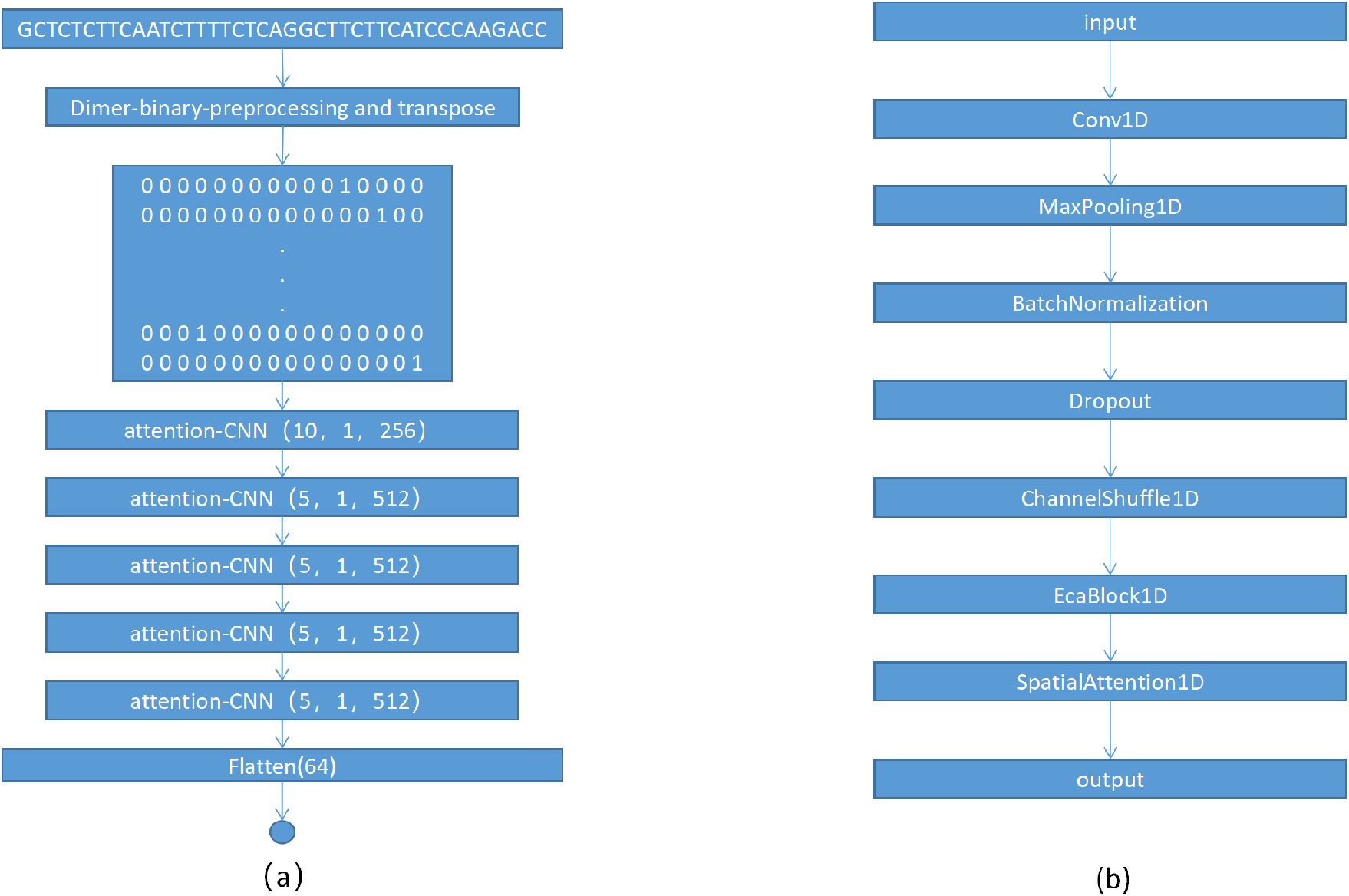
(a) The basic frame diagram of the model. (b) Implementation Details of the attention-CNN Module.

The constant modules in the five attention-CNN modules are the one-dimensional convolution layer, pooling layer, BatchNormalization layer, Dropout layer, and channel shuffle layer. In order to expand the receptive field and learn more features, we set the convolution kernel size of the first convolution layer to 10 and the number of convolution cores to 256. The size of convolution kernels in the next four layers is 5, and the number of convolution kernels is set to 512. Reducing the size of convolution kernels by half is to prevent overfitting while doubling the number of convolution kernels is to offset the insufficient feature learning caused by reducing the size of convolution kernels by half. ReLu is set to activation function:

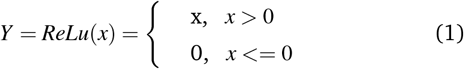

After the convolution core is the pooling layer and the Batch-Normalization layer. We set the pooling size to 2 means that the convolution of each channel in the next layer is reduced to half of the original. The function of the dropout layer is to discard each neuron at a specified probability each time, which is able to discard duplicate cells. We set the probability of the Dropout layer to 0.5, and the number of groups in the channel shuffles layer to 8. For the full connection layer, we set the number of full connection layers in the first layer is set to 128, the activation function to Relu, and the output unit in the second layer to 1. A threshold value greater than 0.5 is predicted as a methylation site, and a threshold value less than 0.5 is predicted as a non-methylation site. Using the sigmoid function as the activation function can ensure that the output is in the interval [0,1]:

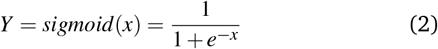

### 2.3 Evaluation metrics

To quantify the performance of ACNN-6mA and compare it with other methods, we used six common performance evaluation indicators: sensitivity (SN), specificity (SP), precision, accuracy (ACC), Matthew correlation coefficient (MCC), F1 score, area under the receiver operating characteristic curve (AUC):

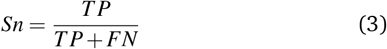

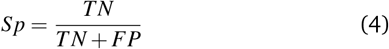

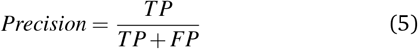

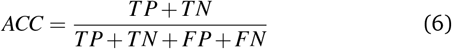

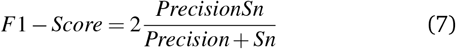

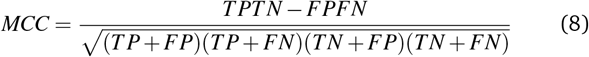

TP, FP, TN, and FN represent the number of true positive, false positive, true negative, and false negative, respectively.

## 3 RESULTS AND DISCUSSION

### 3.1 Pre-training of Rice Dataset

Since there are many studies in the rice dataset, ACNN-6mA is first trained on the rice dataset to obtain the best experimental results and model parameters, and then half of the dataset of other species is used for fine-tuning, and the other half is used as an independent verification set. In other words, a model adapted to this dataset can be obtained. The 6mA-rice-Lv dataset was used for 10-fold cross-validation, and the 6mA-rice-Chen dataset was used for independent verification for independent testing.

Compared with other datasets, the rice dataset has the largest quantity without losing quality, which is conducive to the training of deeper neural networks, while the dataset with a small amount of data is especially easy to fit specific data. We conducted 10-fold cross-validation on our proposed model, SNNRice6mA, Deep6mA, and i6mA-CNN respectively, and found that ACNN-6mA obtained the best experimental results on the 6mA-rice-Lv dataset. MCC, ACC, F1-Score and AUC were 91.29%, 95.64%, 95.67% and 98.90%, respectively. Compared with the Deep6mA model with the best performance, the ACNN-6mA model has improved in five indexes, respectively by 3.11%, 1.60%, 1.49% and 0.72%, as shown in the bar chart in Figure 2(a). It can be found that the results of ACNN-6mA on Sn and Sp are more balanced than those of other models, which is reflected by the relatively high MCC, which also proves that ACNN-6mA has a strong ability to identify positive samples. Figure 3 shows the confusion matrix diagram of the four models on the 6mA-rice-Lv dataset.

**Fig. 2.**
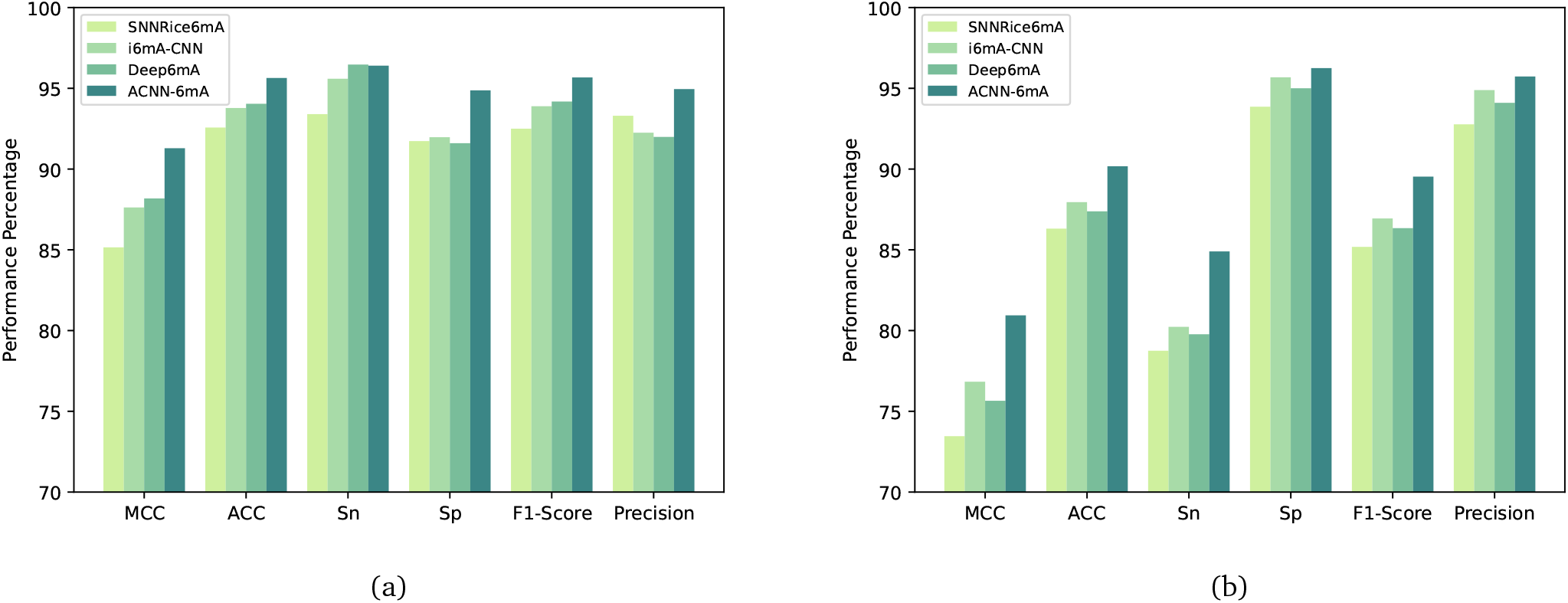
MCC, ACC, Sn, Sp, F1 Core and Precision of SNNRice6mA, i6maCNN, Deep6mA and ACNN-6mA models on rice dataset:(a) 6mA-rice-Lv dataset (b) 6mA-rice-chen dataset

**Fig. 3.**
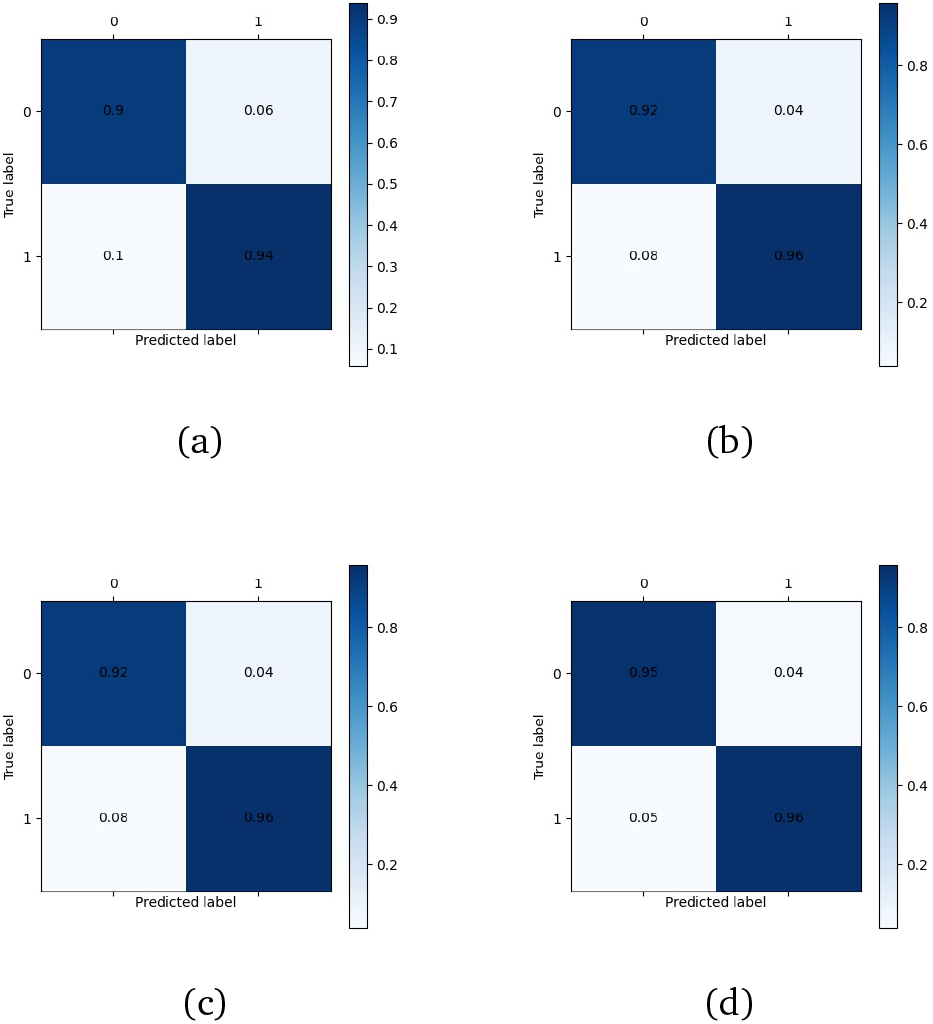
The confusion matrix of SNNRice6mA, i6maCNN, Deep6mA and ACNN-6mA for 6mA-rice-Lv dataset prediction

In order to further study the prediction ability of ACNN-6mA for future rice data, we independently tested the 6mA-rice-Chen dataset with ACNN-6mA, SNNRice6mA, Deep6mA and i6mA-CNN respectively. As shown in Figure 2(b), it can be found that compared with other models, ACNN-6mA has a greater improvement in MCC and ACC values, indicating that ACNN-6mA has a relatively high overall prediction ability for future rice data and a relatively high recognition ability for positive samples. Figure S1 shows the confusion matrix of the four models on the 6mA-rice-Chen dataset. The results show that although the ability of ACNN-6mA to identify positive and negative samples has been significantly improved compared with the previous models, the ability to identify negative samples still needs to be improved compared with the ability to identify positive samples.

Figure 4 shows the ROC curves of ACNN-6mA, SNNRice6mA, Deep6mA and i6mA-CNN on two rice datasets. As shown in Figure S2, we used t-SNE and PCA clustering to analyze the results output from each layer of the rice dataset during prediction and found that after each layer of processing, it can indeed improve the classification effect.

**Fig. 4.**
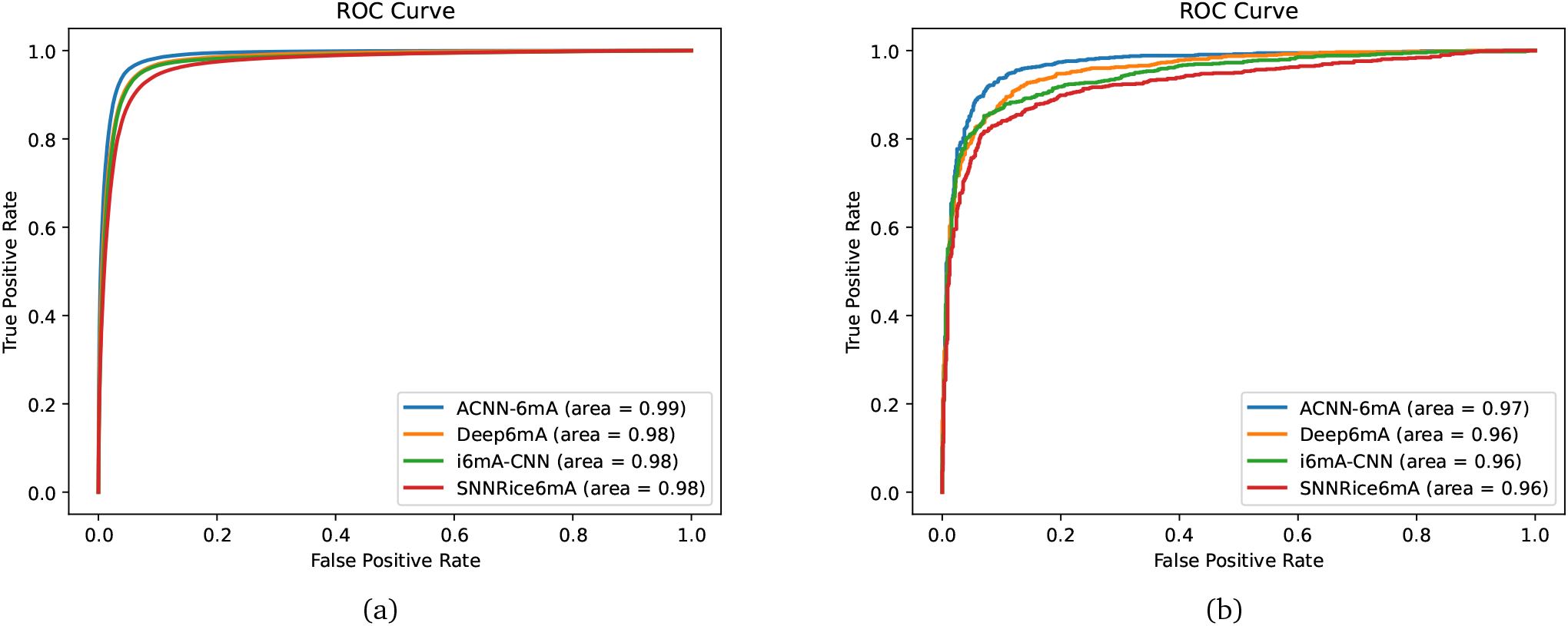
Roc Curve of four models SNNRice6mA, i6maCNN, Deep6mA and ACNN-6mA on rice dataset

### 3.2 Fine-tuning on multi-species dataset

As far as we know, most of the existing m6A models are based on the prediction models obtained from one to four datasets. Developing a model for multi-species prediction is a problem that needs to be solved^49,50^. As far as we know, only iDNA-MS has predicted the m6A sites of a large number of species, but its prediction performance is poor. We selected the higher quality dataset collected and collated by iDNA-MS for model fine-tuning and prediction and found that the performance of ACNN-6mA in most species in its dataset is better than the existing model. Figure 5 and Figure S3 show the MCC, ACC, AUC, SN and SP of ACNN-6mA and iDNA-MS model on 11 species. It can be seen that ACNN-6mA has better prediction performance on 6mA sites than iDNA-MS, and on each species, its AUC, ACC and MCC have been significantly improved.

**Fig. 5.**
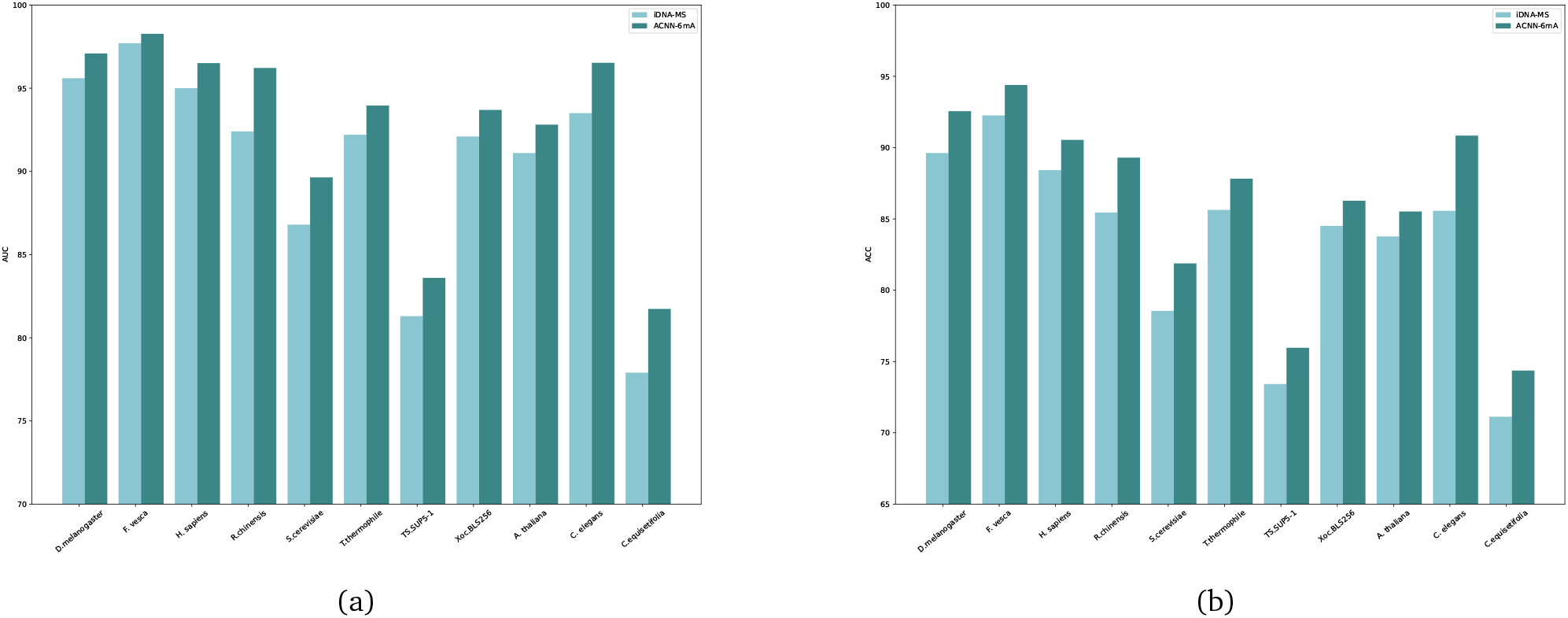
The performance of iDNA-MS and ACNN-6mA on each species independent test set after fine-tuning on D.melanogaster, F. vesca, H.sapiens, R.chinensis, S.cerevisiae, T.thermophile, Tolypocladium, Xoc.BLS256, A. thaliana, C. elegans and C.equisetifolia: (a) AUC (b) ACC

Although the overall recognition ability of ACNN-6mA for methylation sites in the rice dataset is preferable, the recognition ability for negative samples in the rice dataset still needs to be improved. However, through the multi-species training model, we found that ACNN-6mA is not always inferior to the positive sample in identifying negative samples, and although it can achieve elegant results in ACC, MCC, AUC and SP, it is relatively poor in SN for Tolypocladium, F.vesca and C.equisetifolia. Although the performance on other indicators is prominent, the poor performance of SN proves that the ability of ACNN-6mA to identify negative samples needs to be improved. It may be that the ability of ACNN-6mA to identify positive samples makes up for the poor ability to identify negative samples.

We also predicted other species based on the fine-tuning model of each species, and have made some new discoveries. As shown in Figure 6 and Figure S4, the majority of species-trained models can obtain better cross-species prediction results on D.melanogaster, F.vesca, H.sapiens, R.chinensis, S.cerevisiae, TS.SUP5-1, A. thaliana, C. elegans and C.equisetifolia. However, for T.thermophile and Xoc.BLS256 species, most of the species made cross-species predictions or the models trained on these two species on other species are disappointing, which may be due to the large difference in methylation patterns between these two species and the other nine species. For the model of C.equisetifolia, most of the trained models of species have average prediction results, and the trained models can obtain higher accuracy rates for all but T.thermophile, Xoc.BLS256 and Tolypocladium, and even can obtain 95%, 96%, 94%, 93% and 97% AUC values for D.melanogaster, F. vesca, H.sapiens, A.thaliana and C.elegans respectively (although the AUC values for their own datasets are only 91%). For T.thermophile and Xoc.BLS256, the C.equisetifolia-trained model also has high prediction performance. It is very likely that C.equisetifolia can obtain such prominent results in cross-species prediction because its methylation patterns are evenly distributed so that the neural network can fully learn each methylation pattern. For T.thermophile and Xoc.BLS256, not only the prediction performance of models trained on other species is poor, but their mutual prediction performance is the worst (worse than their prediction performance on the other nine species). Therefore, we have reason to believe that T.thermophile and Xoc.BLS256 may contain a small part of the methylation patterns of the other nine species, but the difference between their methylation patterns is also very large.

**Fig. 6.**
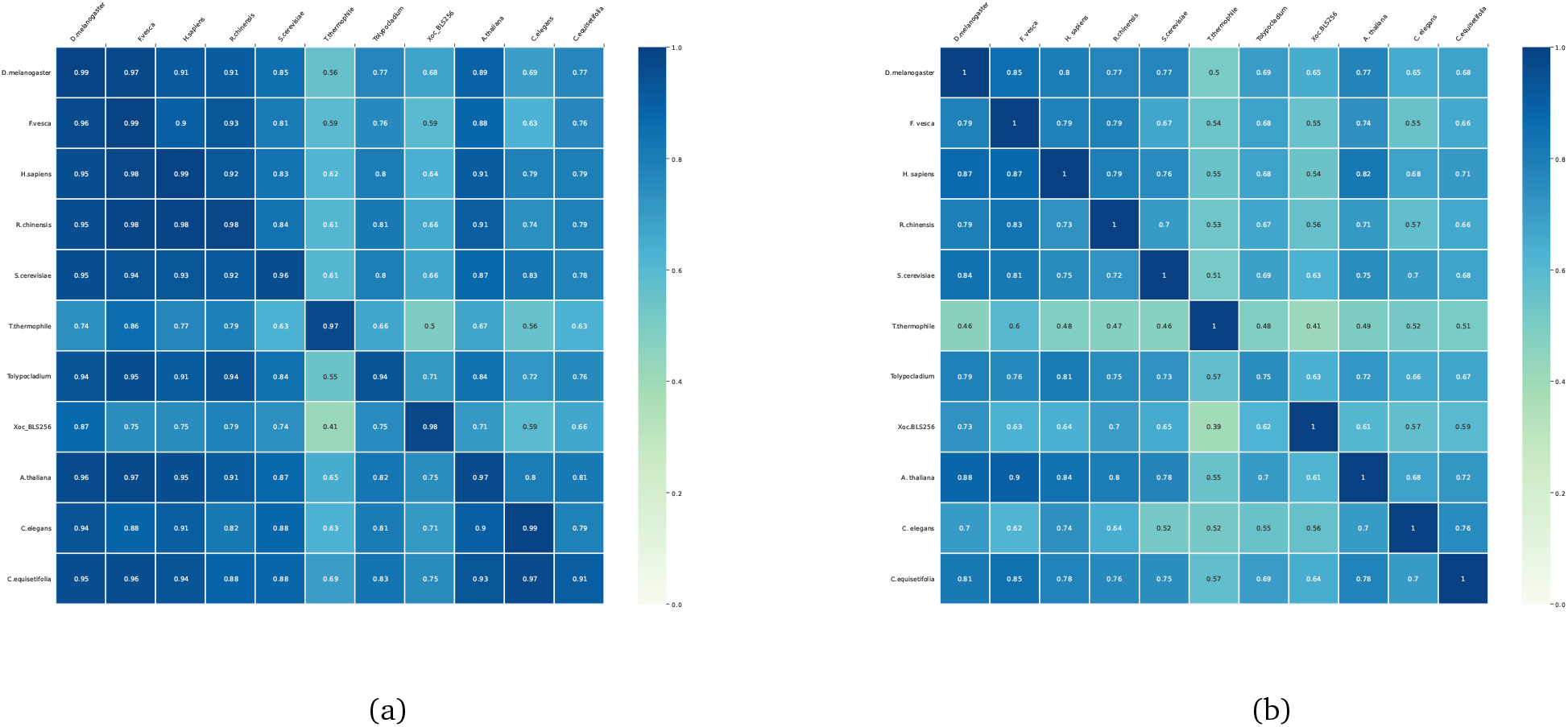
AUC heat maps for predicting 11 species using ACNN-6mA and iDNA-MS models trained on 11 specific species datasets: (a) iDNA MS (b) iDNA MS

We used t-SNE and PCA clustering methods to analyze 11 species respectively and found that D.melanogaster, F.vesca, H.sapiens, R.chinensis, S.cerevisiae, Tolypocladium, A.thaliana, C.elegans and C.equisetifolia have a certain relationship in crossspecies recognition, as shown in Figure 7. We can find from Figure 6 that the species that are more difficult to cluster will not be easy to predict when cross-species prediction is made, but the species that are more difficult to cluster can achieve a higher level of prediction performance for each species when cross-species prediction is made. The mutual prediction performance of D.melanogaster, F.vesca, H.sapiens, R.chinensis, S.cerevisiae, A.thaliana is high, while the prediction performance of these six species for C.elegans and C.equisetifolia is poor. Therefore, we have reason to believe that the difference in methylation patterns among D.melanogaster, F.vesca, H.sapiens, R.chinensis, S.cerevisiae and A. thaliana may not be very large, and the methylation patterns of six species are also an important part of C.elegans and C.equisetifolia methylation patterns. For T.thermophile and Xoc.BLS256, their clustering is difficult and their mutual prediction performance is poor, but their prediction performance for other species is also poor. Therefore, we have reason to believe that their methylation patterns are quite different from those of the other nine species. For each species, we also drew a cluster diagram of its features extracted by the neural network, as shown in Figure 8.

**Fig. 7.**
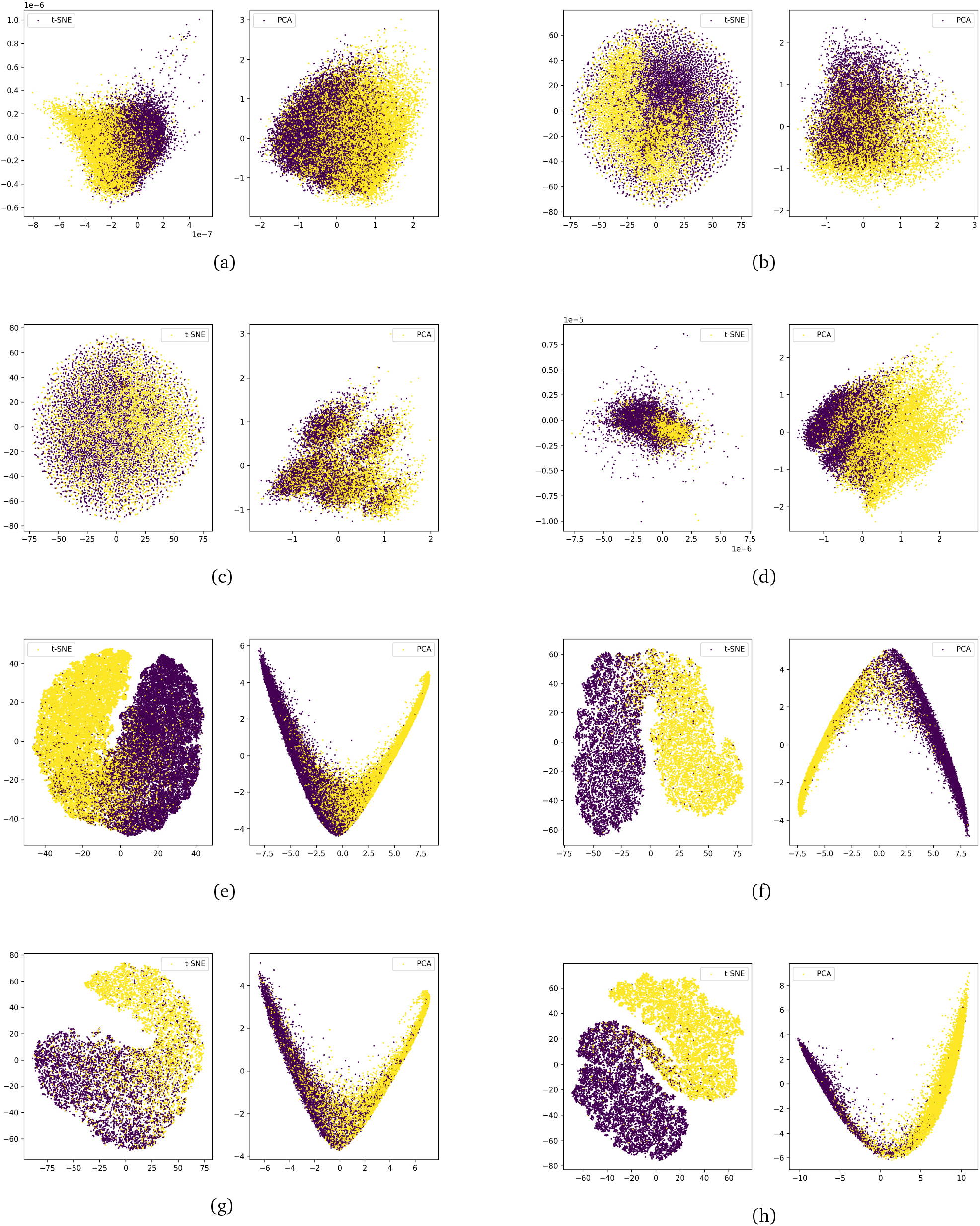
(a), (b), (c) and (d) are respectively the t-SNE and PCA clustering results of A.thaliana, C.elegans, C.equisetifolia and D.melanogaster species on the original data, and (e), (f), (g) and (h) are respectively the t-SNE and PCA clustering results A.thaliana, C.elegans, C.equisetifolia and D.melanogaster species after ACNN-6mA treatment

## 4 Challenges and future work

Although the species we studied include most of the species studied by existing models, compared with the huge methylation biological database, the current popular public dataset is still small, and larger methylation datasets and more balanced data can enable neural networks to learn more features. For the D.melanogaster, F.vesca, H.sapiens, R.chinensis, S.cerevisiae, Tolypocladium, A. thaliana and C. elegans species, the methylation patterns among the species may be similar, so the research based on these eight species can be conducted simultaneously without training the neural network separately. However, the methylation patterns of species C.equisetifolia may be relatively rich, so opening up a larger C.equisetifolia dataset may be very conducive to improving the ability of cross-species prediction. The T.thermophile dataset can achieve high performance on an independent verification set, but its cross-species prediction ability is poor. It may be that there is a large gap between its methylation pattern and other species, and new research may be conducted in the future to make a new cross-species prediction with the species T.thermophile. It may be important to predict the 6mA site of Xoc.BLS256 species, because ACNN-6mA and previous studies have obtained poor performance on this dataset.

## 5 Conclusions

In order to solve the problem of poor performance of machine learning in identifying methylation sites, we first trained a model based on 10-fold cross-validation of the 6mA-rice-Lv dataset and verified it on the 6mA-rice-Chen dataset. The results show that ACNN-6mA can achieve more outstanding results than previous studies on these two datasets. Then we fine-tune based on the data of half of the 11 species pre-trained on the rice dataset, and the obtained model can achieve more pleasurable classification performance on the remaining half of the data of 11 species. We also used the trained models on 11 species to predict methylation sites of other different species across species and achieved excellent results. By analyzing the cross-species pre-training results of 11 species and clustering analysis of the original data, we found some clues about the cross-species training performance of our neural network.

Our model has a strong recognition ability for rice 6mA methylation sites, but its recognition ability for negative samples is stronger than other previous models, but it still needs to be improved. Although ACNN-6mA pre-trained by rice dataset can achieve good performance in multi-species methylation 6mA site recognition, for T.thermophile, although its independent verification set has prominent performance, its cross-species prediction performance is poor. In the future, we may need some new species with similar methylation patterns to study their crossspecies recognition ability. For species C.equisetifolia, although the model trained on its model has strong cross-species recognition ability in most models, its own independent verification effect is poor, so using a larger species C.equisetifolia in the future may achieve superior results in methylation recognition of most species. For Xoc.BLS256 species, we found that the model is not only poor in its own independent verification set but also poor in cross-species prediction. In a word, ACNN-6mA based on rice dataset pre-training can predict future data on the A. thaliana, C. elegans, D.melanogaster, F. vesca, H. sapiens, R.chinensis, S.cerevisiae, T.thermophile and Xoc.BLS256 independent verification set with high performance and greatly improve the A. thaliana, C. elegans, C.equisetifolia, D.melanogaster, F. vesca, H. sapiens, R.chinensis, S.cerevisiae and Tolypocladium cross-species prediction ability compared with previous research.

## Declarations

### Ethical Approval

not applicable

### Authors’ Contributions

Jian Guo Bai is responsible for writing the paper design and program design, Hai Yang is responsible for guiding work

### Availability of data and materials

The code involved in this experiment and the data set collected are available as open-source (https://github.com/jrebai/ACNN-6mA).

