## Supplementary material for "ACNN-6mA Prediction of N6-Methyladenine Loci in Multiple Species Based on Rice Dataset Pre-training Model^†^": Suppliment

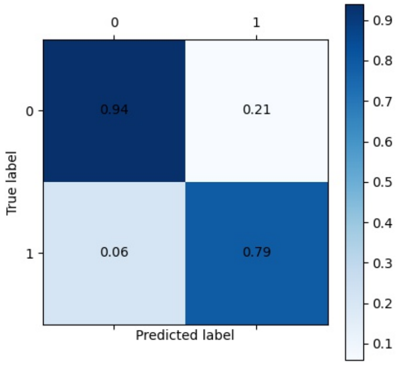


(a)

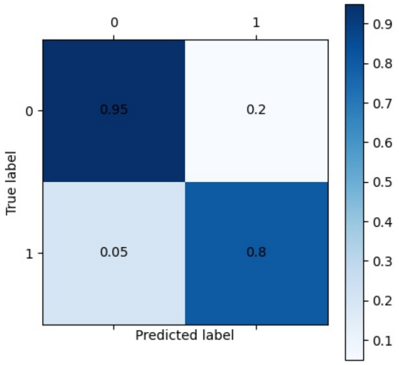


(b)

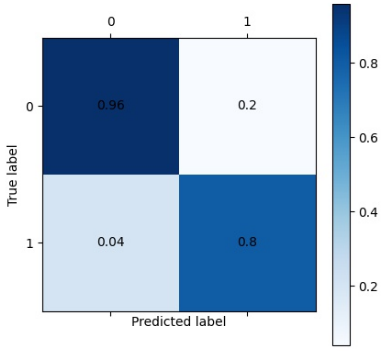


(b)

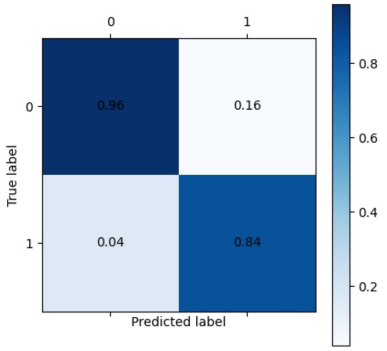


(b)

Figure 1: The confusion matrix of SNNRice6mA, i6maCNN, Deep6mA and ACNN-6mA for 6mA-rice-Chen dataset prediction: (a)SNNRice6mA (b)i6maCNN (c)Deep6mA (d)ACNN-6mA

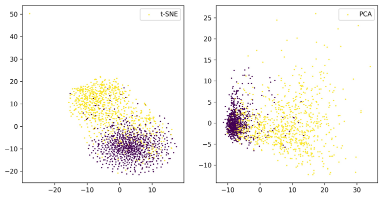


(a)

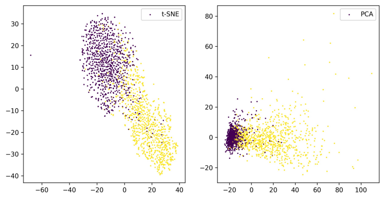


(b)

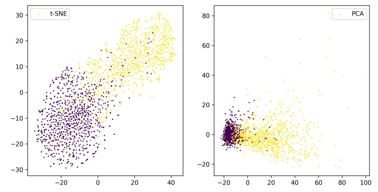


(c)

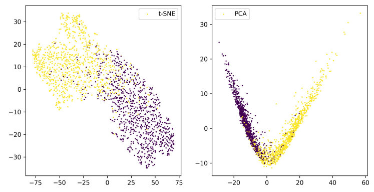


(b)

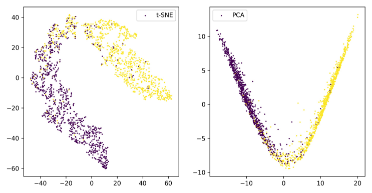


(d)

Figure 2: (a), (b), (c), (d) and (e) are the clustering results of t-SNE and PCA output from the first layer to the fifth layer respectively

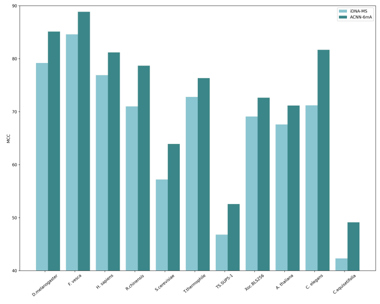


(a)

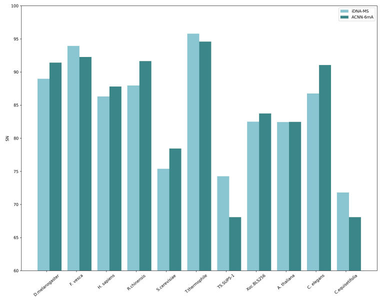


(b)

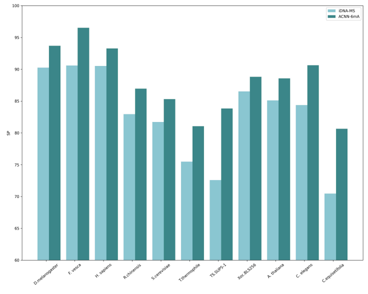


(c)

Figure 3: MCC, Sn, Sp of 11 species on iDNA-MS and ACNN-6mA: (a)MCC (b)Sn (c)Sp

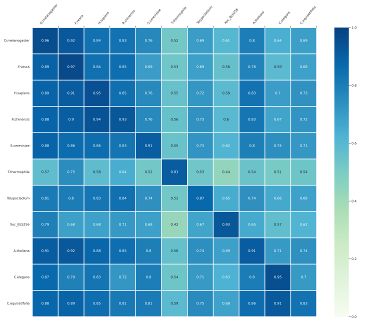

(a)

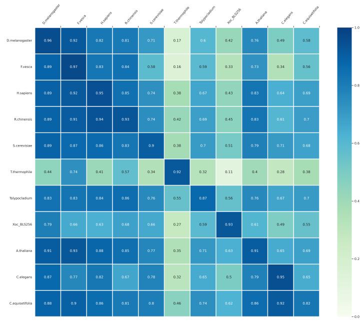

(b)

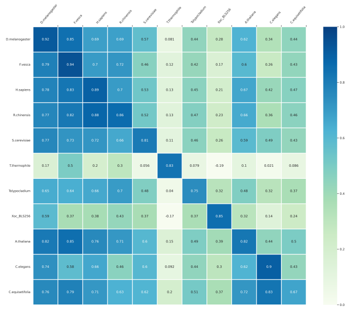

(c)

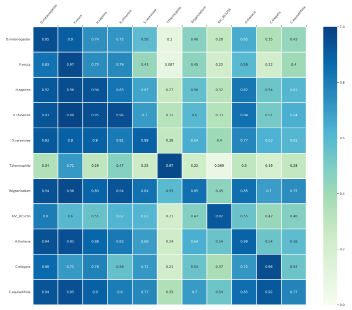

(b)

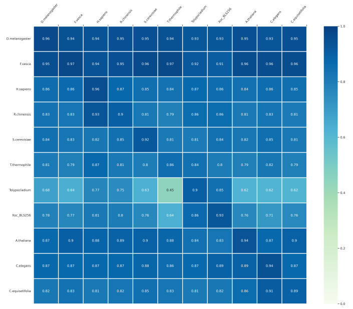

(d)

Figure 4: Results of cross species prediction of ACNN-6mA after fine tuning of 11 species:(a)ACC (b)F1-Score (c)MCC (d)Sn (e)Sp

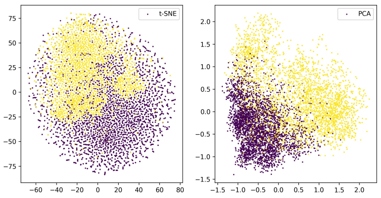


(a)

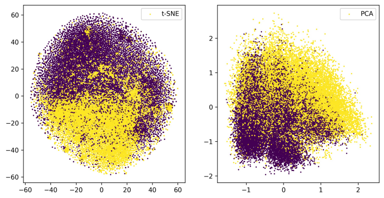


(b)

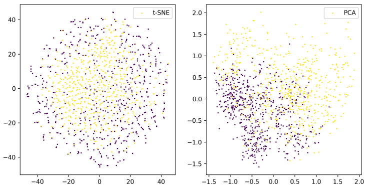


(c)

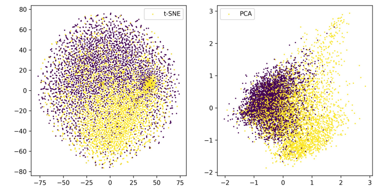


(d)

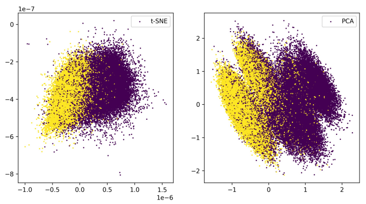


(e)

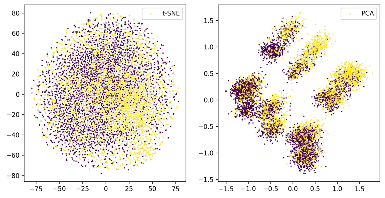


(f)

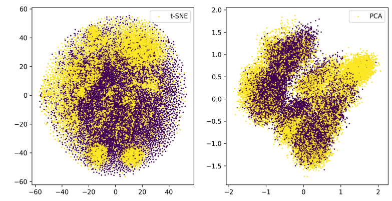


(g)

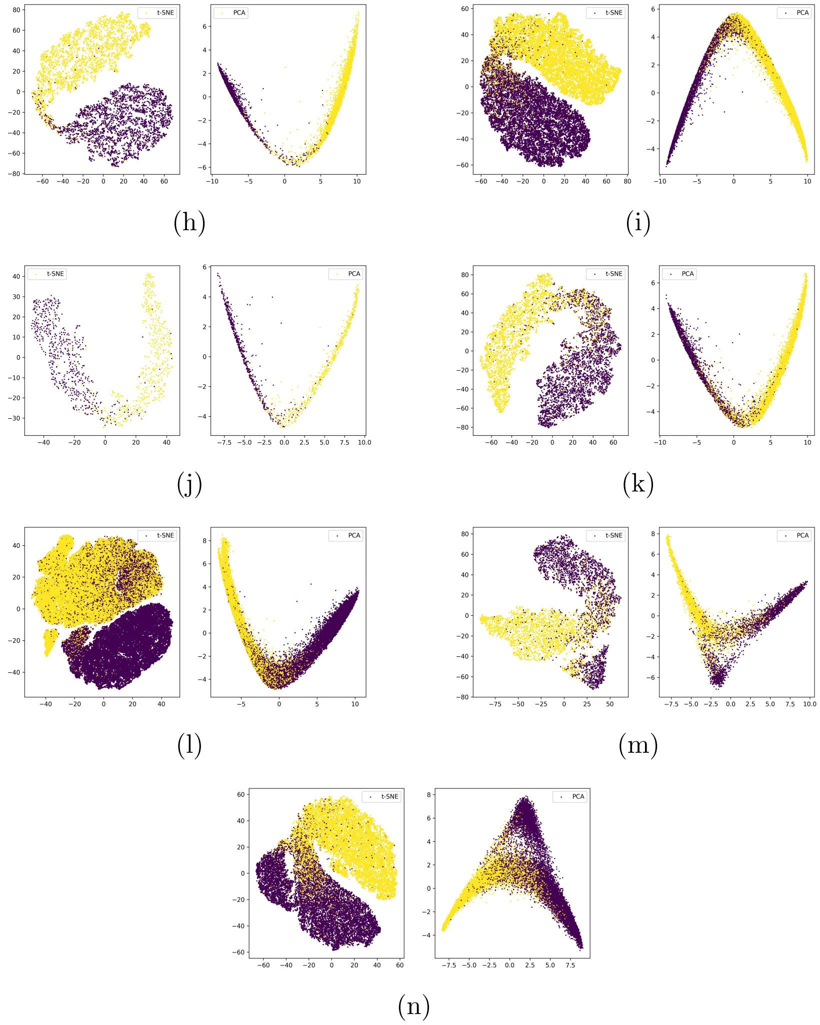


Figure 5: Clustering results of multi-species original data and clustering results after neural network processing, Clustering results of original data: (a)F.vesca (b)H.sapiens (c)R.chinensis (d)S.cerevisiae (e)T.thermophile (f)Tolypocladium (g)Xoc_BLS256. Clustering results after neural network processing:(h)F.vesca (i)H.sapiens (j)R.chinensis (k)S.cerevisiae (l)T.thermophile (m)Tolypocladium (n)Xoc_BLS256.
